# A hydrophobic cluster controls long-range allostery in the TRMT2A RNA recognition motif

**DOI:** 10.1101/2025.11.05.686750

**Authors:** Mohammed Khaled, Lisa Johannknecht, Oscar Palomino-Hernandez

**Author notes:** these authors contributed equally (Mohammed Khaled, Oscar Palomino-Hernandez).

## Abstract

TRMT2A has emerged as a disease-modifying target in polyglutamine (PolyQ) models, yet the conformational preferences and allostery of its RNA recognition motif (RRM) remain poorly resolved. Here we combine extensive atomistic molecular dynamics with Markov state modeling (BHMSM), transition path theory, and structure-based pocket analysis to map the conformational landscape of the human TRMT2A RRM.

We resolve six metastable states and show that a hydrophobic cluster centered on F92–W134–L133 governs their interconversion. We further identify residues contributing to RNA strand recognition and reveal state-specific cryptic pockets consistent with reported TRMT2A RRM small molecule inhibitors.

Together, these results support a hinge–gate model in which a soft, defect-enabled *α*2 segment and a loop 5 hydrophobic cluster coordinate long-range communication between the RNP face and opposite side, yielding testable mutational predictions and state-specific opportunities for allosteric control of TRMT2A in polyQ disease contexts.

## 1 Introduction

Polygluatmine (polyQ) diseases are a subset of trinucleotide repeat expansion disorders. These expansions produce abnormally long polyQ tracts in the corresponding proteins, which adopt *β*-sheet–rich conformations and aggregate into insoluble inclusion bodies, hallmarks of neuronal pathology.[1, 2] Mounting evidence indicates that pathways controlling RNA processing and translation can modulate polyQ toxicity, highlighting RNA–protein interactions as promising therapeutic entry points. In this context, human tRNA methyltransferase 2 homolog A (*h*TRMT2A) — a tRNA m^5^U methyltransferase — has emerged as a potential target for polyQ toxicity.[3] *h*TRMT2A catalyzes the methylation of U_54_ in the tRNA T-loop to form m^5^U_54_, an RNA modification essential for tRNA stability and translational fidelity.[4, 5] Thus, rather than directly engaging with polyQ sequences or their transcripts, inhibition of *h*TRMT2A mitigates polyQ-associated phenotypes by limiting the production of the expanded polyQ proteins.

The RNA binding process of *h*TRMT2A is primarily mediated by its N-terminal RNA recognition motif (RRM). [5, 6] This RRM adopts the canonical *β*1–*α*1–*β*2–*β*3–*α*2–*β*4 fold, as other RRMs.[7] This domain contains two highly conserved short sequence motifs known as RNP1 and RNP2, on the *β*3 and *β*1 motifs, respectively. Aromatic residues within these motifs, such as F106 and F113, engage in *π*–*π* stacking with nucleobases and can intercalate between adjacent bases to stabilize binding.[8, 9] X-ray structures of the *h*TRMT2A RRM (PDB: 7NTO, 7NTN) revealed the atomistic details of this domain and suggested the existence of a cryptic binding site located between the *α*1 helix and loop 5, opposite to the RNP1/2 RNA-binding surface. Furthermore, this work also hypothesized an allosteric communication mechanism connecting the RNP on one side and W134 from loop 5 in the opposite cryptic site, despite their considerable spatial separation.[10]

Cryptic pockets of this type are typically transient and sparsely populated in equilibrium ensembles, making them difficult to assess from static structures alone. However, cryptic pockets also show strong potential as therapeutic targets, offering opportunities to address otherwise *undruggable* proteins.[11, 12] Capturing their formation and coupling requires methods that resolve rare, slow transitions. While classical molecular dynamic simulations (MD simulations) affords atomistic detail, trajectories often undersample millisecond–second events.[13, 14] Markov state modeling (MSM) addresses this limitation by combining multiple trajectories into a single kinetic model that maps metastable basins and their interconversion rates.[15]

Here, we combine extensive atomistic MD simulations with MSM analysis to delineate the conformational landscape of the *h*TRMT2A RRM. We identified six metastable states, of which two dominate the equilibrium population and serve as kinetic hubs. We also uncover robust allosteric communication between the RNP1/2 face and the opposing surface, mediated by a compact hydrophobic cluster. Together, these results support a mechanistic model in which this internal hydrophobic switch can impact on the conformational landscape of the RNP1/2 face, reshaping the RNA-binding surface in a state-dependent manner. By localizing the key residues and state-specific conformations underlying this coupling, our work provides testable hypotheses for mutational validation and highlights actionable sites for allosteric modulation in the context of polyQ disease biology.

## 2 Materials and Methods

### 2.1 Simulation settings of the *h*TRMT2A RRM motif

To comprehensively sample the conformational landscape of *h*TRMT2A, we employed an iterative simulation and clustering protocol. The process was initiated by selecting 50 distinct conformations from a prior replica-exchange simulation ensemble.[10] Each of these structures was then used to seed an independent 1000 ns MD simulations. Following this, the resulting trajectories were subjected to geometric clustering to identify 50 new representative conformations. This entire procedure—running 50 simulations followed by clustering—was repeated two more times, for a total of three cycles. The final iteration yielded a refined set of 150 trajectories for subsequent analysis. For each conformation, the system was prepared as follows: The protonation states of the residues were determined using PROPKA.[16, 17] The system was described using the AMBER ff14SB force field [18], at 4 fs time-step through HMR (hydrogen mass repartitioning).[19] The system was solvated in a dodecahedral box with TIP3P water molecules [20], and JC (Joung-Cheatham) counterions (Cl^−^ and Na^+^) were added to neutralize the system to a concentration of 0.15 mol/L.[21] The systems were prepared prior to the production run through energy minimization, 1 ns NVT equilibration, and 1 ns NPT equilibration. Temperature control was managed with the Bussi-Donadio-Parrinello thermostat [22], while pressure control was done with the Bernetti-Bussi barostat. [23] The SHAKE algorithm was used to constrain bond lengths involving hydrogen atoms.[24] An 8 Å cutoff was used for vdW and electrostatic interactions, with particle mesh Ewald (PME) as the long-range electrostatic decomposition approach.[25] All simulations were performed with GROMACS 2022.[26]

### 2.2 Markov State Models

#### 2.2.1 Feature selection

To identify a feature set that best captures the essential dynamics of the system, we first calculated the centers of mass (COMs) for all residue side chains with PyEMMA (version 2.5.11) [27], resulting in 78 COMs. Pairwise distances between said COMs were then determined, yielding 3003 side chain-distances. To reduce dimensionality and identify the most relevant dynamical motions, time-lagged independent component analysis (TICA)[28] was performed with a lag time of 20 ns, after which 40 dimensions were kept. Correlations between the initial feature set and the TICA components were subsequently computed and used as the basis for feature pruning. The first six independent components (ICs) were selected, as they exhibited the strongest correlations with the features (Supplementary Figures S1 and S2).

To refine the feature set, several filtering criteria were applied. Residue pairs involving N- or C-terminal residues were excluded, as well as direct neighbors (|*i* - *j* | = 1 or 2). In addition, for each residue pair (*i*, *j*), a 3×3 neighborhood consisting of [(*i* ±1, *j* ±1), (*i* ±1, *j*), (*i*, *j* ±1), (*i*, *j*)] was considered, and if multiple neighbors exceeded the threshold, only the pair with the highest correlation was retained. Finally, pairs with a low correlation to the ICs were also excluded. A correlation threshold of 0.75 was applied for IC1, while a threshold of 0.45 was used for IC2 - IC6. After applying these filtering steps, a set of 92 unique residue pairs was obtained. To further capture the structural motions, two additional descriptors were included: the dihedral angle defined by the C*_α_* of residues F86-F96-L129-A119, which characterizes the relative motion between helices *α*1 and *α*2, and the angle between the rings of F92 and W134, which describes aromatic coupling.

#### 2.2.2 Markov State Model

The feature set was then used to construct an initial MSM with six states. TICA was applied with a lag time of 20 ns and 30 dimensions, followed by *k*-means clustering into 1000 microstates. [27, 29] An MSM was built using a lag time of 100 ns. Six metastable states were obtained. However, the implied timescales (ITSs) of this initial model showed high variance. Thus, representative structures of the six metastable states were visualized and analyzed to identify the most kinetically relevant features. From this analysis, a refined set of 18 features was selected, consisting of COM distances (W134-F92, W134-L133, F92-L133, L77-F92, L77-L133, L77-W134 and L77-F86), C*α*-C*α* distances (F86-K104, F86-N79, R95-H130, L129-A132, L129-W134, L133-R137, A132-P138, F86-L105 and F86-L134), the dihedral angle describing helix motion, and the angle between the planes of W92 and W134 (Figure 1B).

**Figure 1:**
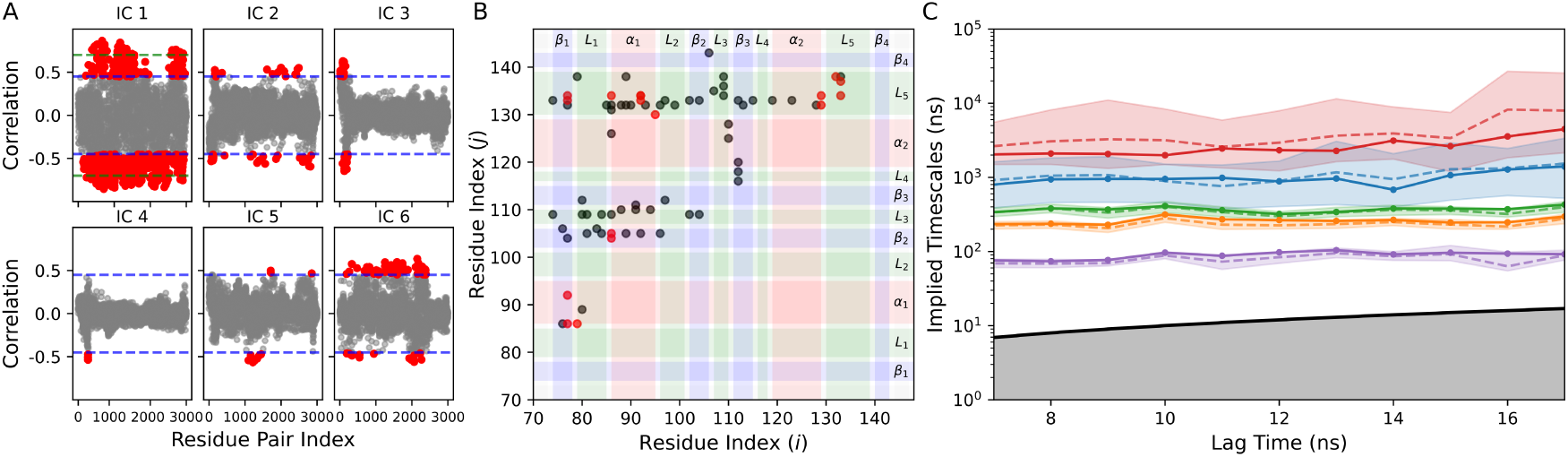
(A) Feature–TICA correlations of residue pairs for the first six independent components, shown as grey dots. Correlation values above 0.45 are shown in red, after the horizontal blue line threshold. The green line threshold indicates correlation values larger than 0.70. (B) Scatter plot of residue pairs. The full feature set of 92 residue pairs is shown in gray, while the 16 residue pairs from the final 18 feature set are shown in red. The protein secondary-structure elements are highlighted as *β*-strands (blue), *α*-helices (red), and loops (green). (C) Implied timescales of the five slowest dynamical processes. Dashed lines represent BHMSM sample means, while solid lines correspond to maximum-likelihood estimates. Shaded areas indicate 90% confidence intervals. The black line with the grey-shaded region denotes processes faster than the lag time.

#### 2.2.3 Hidden Markov State Model

The final feature set was used to build a Bayesian Hidden Markov state model (BHMSM). [27, 29] First, TICA with 2 ns lag time was used to reduce the dimensions to two dimensions. The *k*-means cluster algorithm was used for the discretization in 100 states. The discretized states were used to evaluate the ITS at different numbers of macrostates, where 6 was the maximum number of macrostates that we were able to isolate. Based on this, a lag time of 10 ns was selected for constructing the BHMSM.

The Chapman-Kolmogorov test was employed to validate the Markovianity of the six metastable states by comparing the model-predicted residence probabilities with those from the estimated model. As shown in Figure S3, the predicted probabilities deviate only slightly from the estimated values, thereby confirming the Markovianity of the model.

The state populations, mean state lifetimes, relaxation timescales, and mean first passage times (MFPTs) were calculated after fitting the model samples to a Gaussian distribution using a Gaussian mixture model (GMM) with scikit-learn [30].

#### 2.2.4 Cavity detection

Cavities were identified using MDpocket [31], which is based on the fpocket cavity detection algorithm.[32] Fpocket detects potential binding cavity by generating an ensemble of *α*-spheres that capture the local geometry of the protein. Each *α*-sphere represents a void region tangent to four protein atoms, providing a geometric approximation of the cavity’s shape and size. The cavities were characterized by their volume, calculated from the total number and spatial distribution of *α*-spheres, and by the mean local hydrophobic density, which quantifies the clustering of hydrophobic *α*-spheres within each cavity.

## 3 Results

### 3.1 Feature selection and MSM construction

To capture motions relevant to allosteric communication within the *h*TRMT2A RRM, we first enumerated inter-residue distance features across the domain and quantified their correlation with the slow TICA components (Figure 1A). Among the six slowest ICs, IC 1, 2, 5, and 6 captured collective rearrangements across the helical and *β*-sheet faces, whereas IC 3 and 4 mostly reflected terminal residue fluctuations (Supplementary Figure S2). We therefore retained IC 1, 2, 5, and 6 and pruned the feature set by correlation to these ICs, yielding 92 residue-pair distances (Figure 1B, grey dots). These 92 features are enriched for (i) contacts between loop 5 and the rest of the protein, (ii) couplings between *β*2/*β*3 and *α*1 plus adjacent loops, and (iii) packing between *α*1 and *β*1, foreshadowing a coupling between the helical surface and the RNP sheet.

An initial MSM trained on the 92 features resolved six provisional macrostates. However, the ITSs did not converge across lag times, indicating a lack of ITS independence likely due to the high dimensionality of the system and suboptimal discretization. We thus examined representative structures from each macrostate (lag time 100 ns) and identified key slow motions focused in loop 5 and the *α*1-helix (Figure 1B). This analysis guided a targeted refinement to 18 kinetically informative features that preserved residue-level interpretability, consisting of COM distances (W134-F92, W134-L133, F92-L133, L77-F92, L77-L133, L77-W134 and L77-F86), C*α*-C*α* distances (F86-K104, F86-N79, R95-H130, L129-A132, L129-W134, L133-R137, A132-P138, F86-L105 and F86-L134), a dihedral angle describing the relative motion between helices, and the angle between the planes of W92 and W134 (Figure 1B).

Using this 18-feature set, we trained a BHMSM. The implied timescales (Figure 1C) together with Chapman–Kolmogorov validation (Figure S3) supported the Markov assumption at the chosen lag time, which allowed us to extract kinetic information of this model.

### 3.2 Structural properties of the six metastable states of in *h*TRMT2A

The optimized BHMSM resolved six metastable states (S1–S6) describing the slow dynamics of the *h*TRMT2A RRM. Among these, two states dominate the equilibrium ensemble: S6 (62% occupancy) and S5 (24%), together accounting for approximately 86% of the conformational ensemble (Figure 2A, Supplementary Table S1). Both states closely resemble the crystal structure, exhibiting the lowest heavy-atom RMSD values. The remaining four states (S1–S4) are sparsely populated, each contributing 2–5% of the total equilibrium probability. To elucidate the structural basis of these differences, we compared the secondary-structure composition, residue interactions involving loop 5 and the *α*2-helix, the relative orientation of the *α*1/*α*2-helices, and local flexibility across all six states (Figure 2B–D).

**Figure 2:**
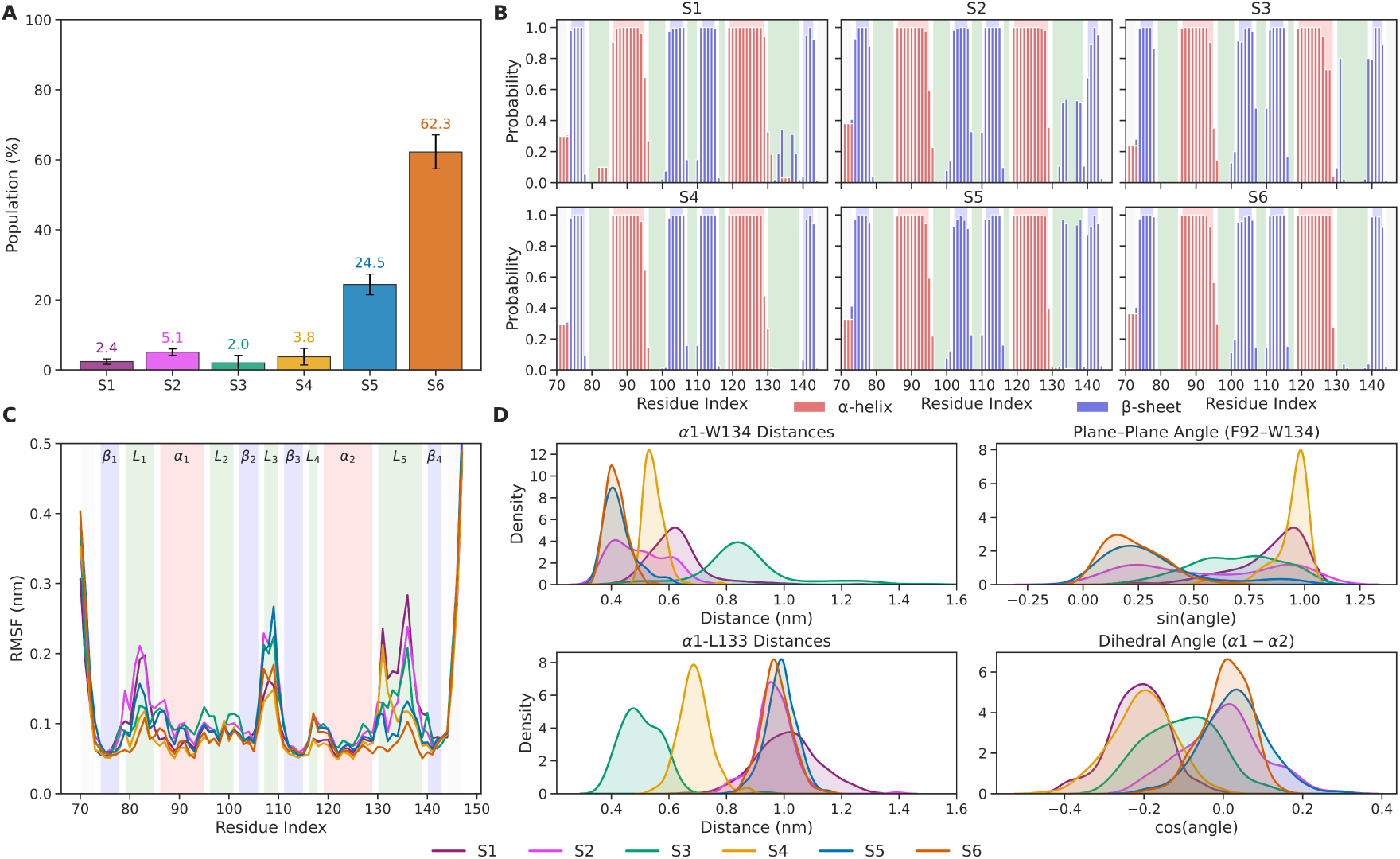
(A) Stationary populations of the metastable states. Error bars represent the standard deviation. (B) Secondary structure probabilities per residue for the metastable states. Bars indicate the cumulative probabilities of *α*-helix (blue) and *β*-sheet (red) content, the remaining fraction up to 1.0 corresponds to random coil. (C) C*α* rootmean-square fluctuations (RMSFs) of the metastable states. RMSFs were calculated after alignment to the crystal structure. The RMSFs of the metastable states are shown in sky blue, pink, green, orange, blue, and red, respectively. The protein secondarystructure elements are highlighted as *β*-strands (blue), *α*-helices (red), and loops (green). (D) Distributions of structural features for the metastable states: The side-chain COM distances between F92-W134 (top, left), the angle between the planes of W92 and W134 (top, right), the side-chain COM distances between F92-W133 (bottom, left) and the dihedral angle describing helix motion (bottom, right).

The metastable states show notable differences in their secondary-structure content (Figure 2B). Taking the most populated state, S6, as a reference, S5 exhibits additional *β*-structure within loop 5 (residues 133–134 and 137–138), consistent with a slightly more rigid conformation. This observation suggests that S5 and S6 interconvert through localized *β*-sheet formation in loop 5. The third most abundant state, S2, shows a similar but less pronounced rigidization in this region, indicating partial stabilization of the same structural motif.

The least populated states (S1, S3, S4) display broader rearrangements involving both the *α*2-helix and *β*-sheet elements. In S4, *β*-sheet structure is partially lost at residues 78 (*β*1), 100 (*β*2), and 116 (*β*3), accompanied by a slight extension of the *α*2-helix near residue 130. S1 shows a comparable pattern of *β*-sheet loss but with the additional ordering within loop 5 as described before. The *α*2-helix content in S1 also extends through residues 130–131. Conversely, S3 gains *β*-sheet content at residues 107 (*β*2), 110 (*β*3), 131 (loop 5), and 139 (*β*4), while its *α*2-helical segment shortens, losing helical propensity at residue 129. Overall, these patterns indicate that loop 5 and the C-terminal portion of the *α*2-helix are key structural hotspots modulating the conformational heterogeneity of the *h*TRMT2A RRM.

The root-mean-square fluctuations (RMSF) (Figure 2C) show that deviations from the crystal structure are most pronounced in loop 1, loop 3, and loop 5. States S6 and S5 are the most rigid, whereas the remaining states display elevated flexibility in these regions. Notably, residues 132–134 within loop 5 do not exhibit the same increase in mobility as neighboring residues, suggesting the presence of local stabilizing interactions that anchor this segment.

The gain of secondary structure within loop 5, together with its localized rigidity at residues 132–134, prompted us to investigate the interactions responsible for this stabilization. In the most populated metastable states, S6 and S5, W134 remains buried in close contact with F92 from the *α*1-helix, forming a nearly parallel *π*–*π* stacking arrangement (Figure 1D, upper row). Given this characteristic, we thus refer to the F92/W134 as an aromatic lock. This lock, quantified through both inter-residue distances and ring-plane orientation angles, keeps a surrounding compact hydrophobic core that maintains a rigid conformation on the *h*TRMT2A RRM. In S2, the aromatic lock becomes slightly disrupted due to the formation of the transient *β*-sheet content within loop 5, as reflected in the increased inter-residue distances and a wider distribution of plane orientations. This intermediate state thus alternates between mixed *π*–*π* and hydrophobic interactions.

Less populated states exhibit distinct rearrangements of this aromatic lock. In S4, W134 tilts away from F92, adopting a T-shaped ring orientation. Concomitantly, L133 moves inward toward F92, likely compensating for the change of aromatic stacking by shielding the hydrophobic core from solvent exposure. In S3, the shift becomes complete: W134 disengages entirely, and L133 replaces it, effectively redefining this interaction as purely hydrophobic (Figure 1D, lower left). Conversely, in S1, W134 drifts away from F92 without replacement by nearby residues, leaving the site exposed and highly flexible, consistent with its low population (Figure 1D, lower right). In all these states, we observed the formation of a small cavity in place of the broken aromatic lock, depending on the degree of displacement of W134, as suggested by Margreiter et. al. [10].

The dihedral angle between the *α*1- and *α*2-helices mirrors the hydrophobic rearrangements described above. In the closed states S6 and S5, the helices adopt a nearly perpendicular orientation that maintains a compact fold, with S2 retains a comparable geometry with minor deviations. In contrast, S3, S4, and S1 exhibit a shift of the *α*2-helix accompanied by the concerted burial of residue F86, which is solvent-exposed in the closed conformations. This rearrangement not only extends the hydrophobic network toward the domain core but also increases the distance between the RNP region and the helices, creating a secondary cavity, only present in these high energy states.

Together, these observations indicate that the aromatic and hydrophobic interactions within loop 5—particularly those involving W134, F92, and L133—act as a molecular switch that stabilizes the inter-helical orientation. The dynamics of this aromatic lock control both the formation of cavities within the structure as well as the solvent exposure of F86. F86 thus emerges as a reporter residue nearby the RNP1/2 *β*-sheet region, reflecting the state of the aromatic lock and linking conformational changes in loop 5 to alterations at the RNA-binding surface.

### 3.3 Macrostates on *h*TRMT2A seem to be stabilized through defect formation in the *α*2-helix

The analysis above established that the motion of residues within loop 5 plays a key role in shaping the conformational landscape. However, this loop does not operate in isolation: its conformational changes are tightly coupled to the structural plasticity of the adjacent *α*2-helix, which can partially unfold to provide the mechanical leverage required for rearrangement of the hydrophobic cluster.

In this way, the *α*2-helix adopts three discrete conformational states (Figure 3A). The most populated form, designated *α*2-H1 and observed in macrostates S6, S5, and S2, extends the helix to residue 130. Transitions to higher-energy conformations involve either loss of residue 130 from the helical register (*α*2-H2; S3 and S4) or incorporation of residue 131 into the helix (*α*2-H3; S1 and S4). These assignments are corroborated by changes in local backbone angles and diagnostic inter-residue distances (Figure 3B). Within a soft-matter perspective, the partial unwinding of *α*2-which modulates loop 5 motion—constitutes a *connectivity defect*: a local break in the continuity of an otherwise ordered elastic element.[33–36] A useful reference state is a perfectly formed *α*2-helix with a continuous *i* →*i* +4 hydrogen-bond network, expected to occupy a deep, narrow energy well separated from alternative conformations by high activation barriers. Indeed, AlphaFold2 predictions indicate that residues 119–134 have strong intrinsic helical propensity (Supplementary Figure S5). Yet if fully helical in the protein context, the system would become overly rigid and unable to execute the conformational transitions required for allosteric communication.

**Figure 3:**
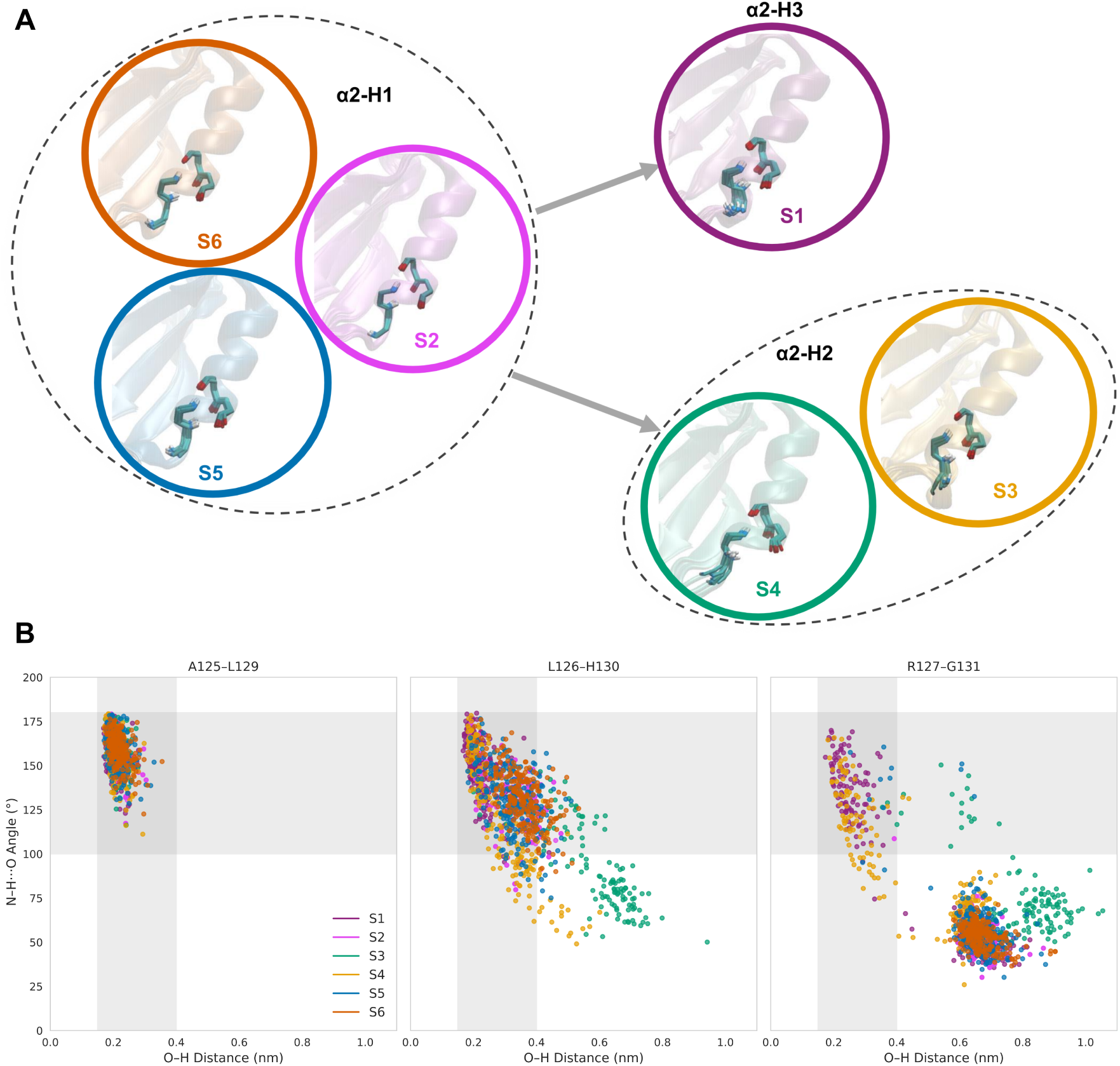
(A) Three conformational states of the *α*2-helix (*α*2-H1, *α*2-H2, and *α*2-H3). The protein is shown in cartoon representation. Hydrogen-bonding atoms of residues A125-L129, L126-H130, and R127-G131 within the *α*2-helix are displayed as sticks. (B) Scatter plots of H, O distances and N-H, O angles describing hydrogen-bond formation between the NH group and backbone oxygen atoms of residues A125-L129, L126-H130, and R127-G131 across the metastable states. The gray-shaded region highlights the geometric criteria for hydrogen bonds.

This finely tuned structural plasticity of *α*2, together with loop 5 motion, defines a coupled hinge–gate mechanism. Partial unfolding of *α*2 modulates loop 5 positioning and thereby tunes accessibility of the cavity adjacent to the RNP face. Thus, the connectivity defect in *α*2 is not a passive mechanical flaw but an active, tunable control element that couples local instability to global functional adaptability. The interplay between these elements underlies the protein allosteric regulation and contributes to its overall thermodynamic stability.

### 3.4 Cavity opening in the *h*TRMT2A RRM is driven by reorganization of aromatic residues W134 or F86

The structural rearrangement surrounding the RNP region in the high-energy conformations facilitates the formation of cavities in the *h*TRMT2A RRM. Two cavities were initially observed: one in the opening of the aromatic lock due to exposure of W134, and another one due to burial of residue F86, near the ribonucleoprotein (RNP). To evaluate other possible cavities and characterize them, we employed MDpocket.[31, 32] Through this approach, we were able to identify and characterize three major cavities: (i) near the aromatic lock F92/W134, (ii) adjacent to F86, and (iii) along the RNP face (Figure 4A). We evaluated these cavities based on their volumes and mean local hydrophobic density; parameters that have been shown to correlate strongly with pocket druggability [37].

**Figure 4:**
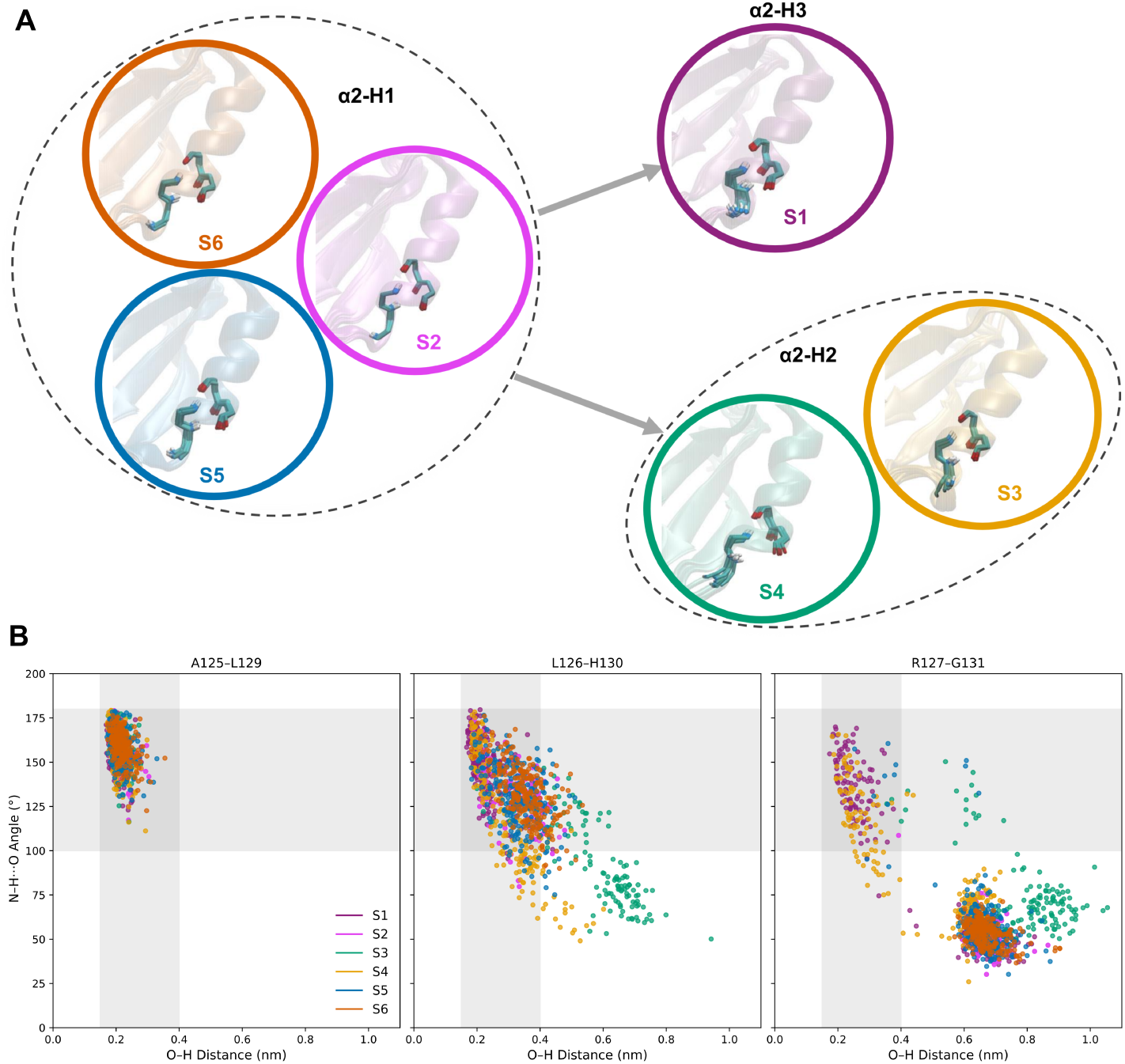
(A) Cavities identified using MDpocket. Cavities are shown as surfaces in green (Cavity 1), yellow (Cavity 2), and purple (Cavity 3). Residues defining each cavity are displayed as sticks in the corresponding cavity color, and the protein is shown in cartoon representation. (B) Relative changes in cavity volume (with respect to S6) and mean local hydrophobic density across the metastable states.

The first cavity, located near the aromatic lock and primarily involving residues from the *α*1-helix, loops 1 and 5, includes H83-S85 (loop 1), S87-V89, R91, F92, R95 (*α*1-hleix), and A132-K135 (loop 5). This cavity exhibits slightly larger volumes in the open conformations (S1 and S4) compared to the closed ones (Figure 4B). Notably, the mean local hydrophobic density is also higher in the open conformations, reflecting the outward movement of W134 away from F92, which exposes part of the hydrophobic core to the solvent and contributes to cavity expansion.

The second cavity, located adjacent to F86, primarily involves residues from the *α*1-helix (F86, V89, R90, L93), *β*2-strand (T103, L104), and *β*3-strand (A112, V114, F116). This cavity is detected exclusively in the open conformations (S1 and S4). Its formation is driven by a shift of the *α*2-helix, burial of residue F86, and an increased separation between the RNP region and surrounding helices. This structural change is evident from the increased distance between S86 in the *α*1-helix and F106 in the *β*2-strand, from 1.022±0.081 nm in the closed conformations (S6 and S5) to 1.205±0.057 nm in the open conformations (S1 and S4), which collectively facilitate cavity formation. Both the cavity volume and mean local hydrophobic density (Figure 4B) are significantly higher in the open states, indicating enhanced cavity accessibility and hydrophobic character. The third cavity is located on the RNA binding surface, defined primarily by the *β*-sheet region. This area is enriched in aromatic residues, particularly F106 and F113, which form *π*-stacking interactions with RNA [8, 9], and also includes C111, which has been experimentally shown to participate in RNA binding. [38] Interestingly, a slight decrease in local hydrophobic density is observed in the open conformations (S1 and S4), coinciding with the inward movement of F86.

### 3.5 Microsecond-scale transitions underlie cavity formation in the *h*TRMT2A RRM

Having established a hinge–gate mechanism in which loop 5 motion is coupled to partial unfolding of the *α*2-helix, which then leads to the formation of the cavities in RRM *h*TRMT2A, we asked *how* the hydrophobic residues—W134, L133, F92, and F86—lead the transition sequence between said macrostates. For this, we applied transition path theory (TPT)[39, 40] to the BHMSM. TPT yielded the dominant transition pathways among the metastable states (Figure 5A, Table 1), as well as state lifetimes (Figure 5B) and slowest relaxation timescales (Figure 5C).

**Figure 5:**
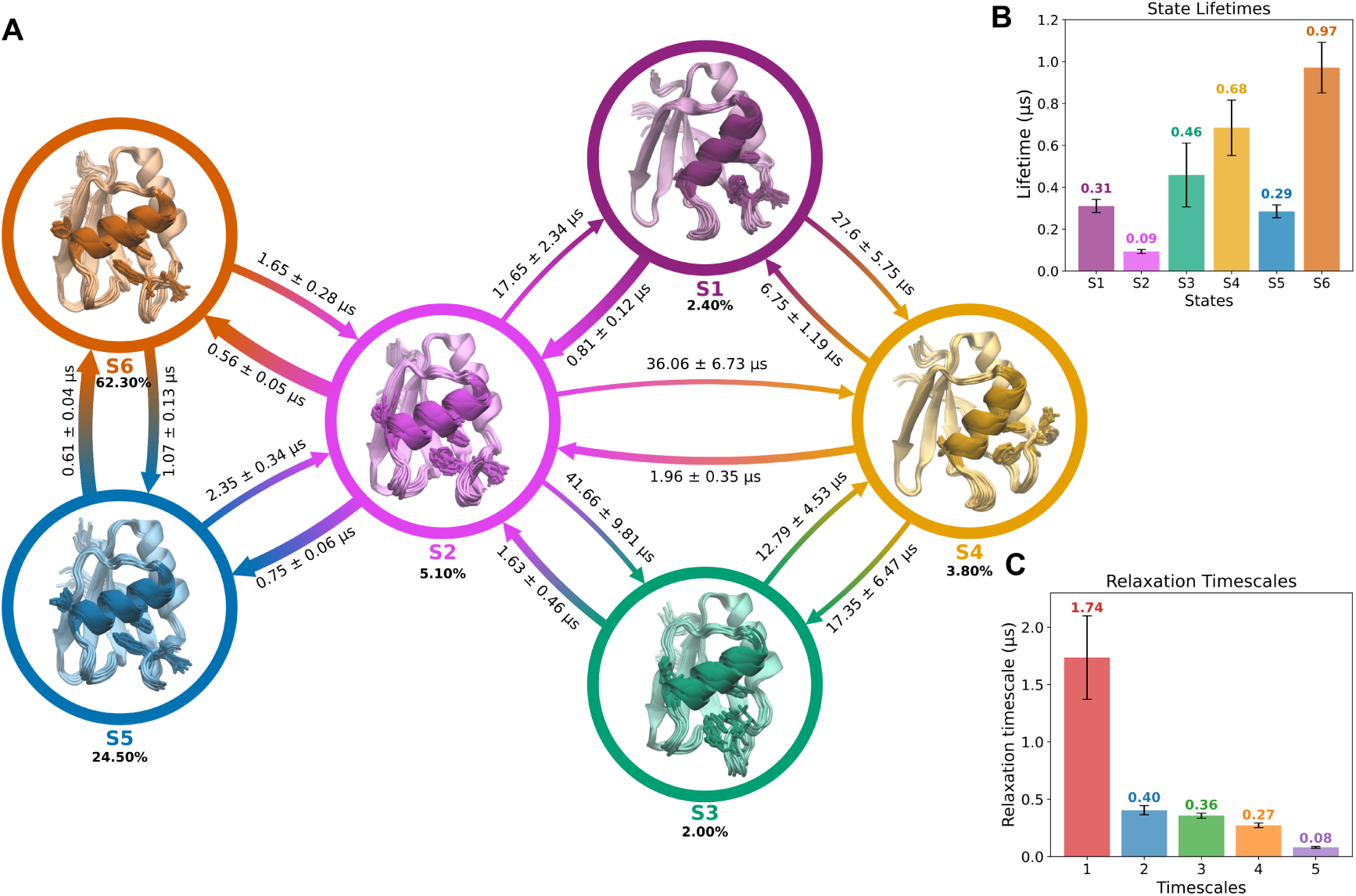
(A) Network diagram of the metastable states and their stationary populations. States are represented as circles, and their populations are shown. Arrows indicate transition rates, annotated with mean first-passage times and their standard deviations. Representative conformations of each metastable state are shown as cartoons, with residues F86, F92, W133, and W134 highlighted as sticks. *α*1-helix is also highlighted in the cartoon depiction. (B) Lifetimes of the metastable states. (C) Slowest relaxation timescales of the model. For both plots, error bars denote standard deviations.

**Table 1:** Transition pathways (Up to ∼90% of the total flux).

| Paths | Percentage (%) |
| --- | --- |
| S6 → S5 → S2 → S1 | 67.12 |
| S6 → S2 → S1 | 20.21 |
| S6 → S2 → S3 → S4 → S1 | 5.12 |

Mean first-passage time (MFPT) analysis revealed that transitions between metastable states span hundreds of nanoseconds to tens of microseconds. State S2 emerges as the principal intermediate bridging the closed F92/W134 core (S6/S5) and the open one (S1/S3/S4). Accordingly, the transition network can be organized into four classes: (i) interconversion within the closed ensemble (S6↔S5), (ii) exchange between S6/S5 and the hub state S2, (iii) transitions between S2 and the open states (S1, S3, S4), and (iv) interconversion among the open states.

Delving further into the transition network: (i) First, transitions between the lowest-energy states S6 and S5 are relatively fast, consistent with rapid formation/disruption of the *β*-structure within loop 5. (ii) Moving forward, the exchange between S6/S5 and S2 involves weakening of the F92–W134 aromatic lock. These processes occur on the order of microseconds and are readily reversible. Across S6, S5, and S2, no large-scale rearrangements are evident, as reflected by minimal changes in the *α*1-helix tilt and the solvent exposure of F86 (Supplementary Figure S4A). (iii) In contrast, transitions from S2 to the open-lock states are markedly slower, reflecting large-scale structural reorganization. The fastest is S2→S1 (∼17 *µ*s), characterized by opening of the F92–W134 lock, tilting of the *α*1-helix, and burial of F86. Paths to S3 or S4 from S2 are slower (∼36–42 *µ*s) and involve partial or complete *α*1 tilting with a corresponding partial/complete burial of F86. The slowest step is S2→S3, which only rewires the lock, substituting W134 with L133, and yields only partial burial of F86. (iv) Finally, interconversion among the high-energy states proceeds via S4, with MFPTs ranging from ∼6 to 28 *µ*s.

Consistent with this network view, S2 is the shortest-lived state (Figure 5B), rapidly giving place either to the closed-lock states (S6/S5) or to the open ones (S1/S3/S4). Within the closed-lock ensemble, S5 is also short-lived and quickly returns to S6 as loop 5 *β*-structure melts. Among the open states, S4 is the longest-lived.

Transition-path theory (TPT) further shows that flux from the most stable closed-lock state (S6) to the most energetic open state (S1) is concentrated in two dominant pathways that account for 87% of the total flux. The major pathway (67%) follows S6→S5→S2→S1: loop 5 first forms *β*-structure (S5), which then destabilizes the aromatic lock, allowing *α*1 tilting, and promoting the burial of F86 (S2), culminating in S1. A secondary pathway (20%) bypasses S5, proceeding S6→S2→S1, indicating that loop 5 *β*-structure formation is not strictly required for opening, even if it stabilizes the preferred multistep route.

In summary, the kinetic analysis highlights a coordinated mechanism with a clear timescale separation: sub- to few-microsecond local rearrangements within the closed ensemble versus tens-of-microseconds transitions that remodel the hydrophobic network and give rise to diverse cavity formation.

## 4 Discussion

We investigated the conformational dynamics of the *h*TRMT2A RRM using atomistic MD simulations. MD simulations typically requires a longer time to converge, undersampling interesting conformations such as higher-energy metastable states and transient (cryptic) pockets. To address this sampling gap, we employed Markov state models (MSMs), which stitch together many trajectories to reconstruct long-timescale kinetics and expose rare events within a single framework.[13]

As outlined in the Introduction, hydrophobic and aromatic residues are indispensable to RRM function, especially for nucleic-acid recognition. Our results highlight a central role for loop 5 in regulating conformational switching among *h*TRMT2A RRM states. In low-energy metastable states S6 and S5, the aromatic side chain of W134 orientates inward toward the *α*1 helix and stacks with F92, minimizing solvent exposure. This *π*–*π* interaction forms an *aromatic lock* that stabilizes a compact ensemble, consistent with the reduced solvent-accessible surface area (SASA) of these states (Supplementary Fig. S4). In higher-energy states (S1, S3, S4), disruption of the F92–W134 interaction breaks the lock, allowing loop 5 to move and enlarging cavity 1. Concomitantly, other residues become more solvent-exposed (Supplementary Fig. S4). This transition is coupled to an outward displacement of the *α*1 helix, which moves away from the core to accommodate an expansion of cavity 2 and the burial of F86.

Loop flexibility depends strongly on sequence composition. Loop 5 comprises A132-L133-W134-R135-G136-R137-P138-L139. While leucine and arginine are common in dynamic loops, the presence of a bulky, hydrophobic tryptophan at position 134 is notable, as tryptophan is underrepresented in loop regions.[41] Nevertheless, W134 (or a phenylalanine at the equivalent position) is conserved across TRMT2A homologs, underscoring its functional importance—a conclusion that aligns with our locking mechanism.

State-dependent cavity formation points to plausible interaction sites for molecular partners. Prior work suggested that motion of W134 can reveal a cryptic pocket.[10] Our MSM indicates that the opening of cavity 1 occurs on the order of tens of microseconds. A second site (cavity 2) emerges from the inward movement of *α*1 and the burial of F86. Among the observed conformational states, S1 and S4—where the F92–W134 lock is broken without compensatory packing and F86 is buried—present the most promising geometries for ligand interaction at said cavities.

The exposure of aromatic side chains is a key determinant of RNA recognition by RRMs.[8] In the crystallized structures of the *h*TRMT2A RRM, two phenylalanines reside on RNP1, but only F113 consistently presents its side chain to solvent; F116 remains buried in the hydrophobic core. On RNP2, F73 is likewise buried, which likely limits its direct role in base stacking. These observations suggest that additional residues may be needed to stabilize nucleobases during binding. In this context, we underscore a potential role for F86 in RNA recognition. In the most stable states (S6, S5, S2), F86 is solvent-exposed. Transitions to conformations in which F86 becomes buried require partial unwinding of the *α*2 helix and a register shift of *α*1—an energetically costly move. We therefore hypothesize that solvent exposure of F86 facilitates initial RNA capture, whereas its burial in higher-energy states constitutes a self-inhibitory switch. Consistent with this model, the volume and hydrophobicity of cavity 3 decrease when F86 is internalized, reducing the likelihood of accommodating nucleobases.

Finally, our model yields concrete, testable predictions for experimental validation: (i) *Lock disruption:* Mutations such as W134A/F and F92A should destabilize S6/S5, shifting populations toward open states; L133A/V may also suppress the S3 “alternative lock.” (ii) *Helix propensity tuning:* Helix-favoring substitutions at 129–131 should bias toward closed states and reduce pocket opening, whereas helix-breaking substitutions (e.g., Pro/Gly at 129–131) should favor opening. These effects can be probed via RNA-binding assays (ITC/SPR) and dynamics-sensitive spectroscopy (NMR CPMG/CEST, smFRET), testing whether reduced binding correlates with stabilized open states (and vice versa).

## 5 Conclusions

By combining atomistic MD simulations with MSMs, we resolve the conformational landscape and allosteric coupling of the *h*TRMT2A RRM. Our results support a coherent mechanism in which a hydrophobic cluster centered on L133/W134–F92, mechanically coupled to a soft, defect-like segment in *α*2, controls the position of *α*1, the exposure of F86, and the breathing of loop 5. This hinge–gate architecture toggles between compact (locked) and open ensembles and remodels solvent exposure across the RNA-binding surface.

Our analysis focuses on the RNA-free RRM. Model choices (force field, water model, protonation), feature selection/discretization, and the absence of post-translational modifications and the C-terminal catalytic domain may influence kinetics and state populations. Extending the MSM to RNA-bound simulations and to full-length *h*TRMT2A will be important to test how RNA engagement feeds back onto the hinge–gate.

The state-resolved pockets and switching residues provide concrete entry points for ligand discovery. In particular, *open-state stabilizers* that wedge near loop 5/*α*1, should disrupt the F92–W134 lock, or stabilize the buried state of F86, thus modulating TRMT2A activity in polyQ disease biology.

In sum, *h*TRMT2A rrm leverages a connectivity-defect–enabled hinge in *α*2 and a hydrophobic microswitch in loop 5 to coordinate distant surfaces and regulate cryptic-pocket opening. This mechanism explains how local instability produces global adaptability and offers actionable targets for allosteric control in polyQ contexts.

## 5 Acknowledgements

This project was funded by the Deutsche Forschungsgemeinschaft (DFG, German Research Foundation – SFB1552 – 465145163”). The authors gratefully acknowledge the computing time granted on the supercomputer MOGON 2 at Johannes Gutenberg University Mainz (hpc.uni-mainz.de). The authors also thank Prof. Dr. Paul Czodrowski and Prof. Dr. Sebastian Seiffert for reading this work.

## Acronyms

BHMSM: Bayesian Hidden Markov state model
COMs: centers of mass
*h*TRMT2A: human tRNA methyltransferase 2 homolog A
ICs: independent components
ITSs: implied timescales
MD simulations: molecular dynamic simulations
MFPTs: mean first passage times
MSM: Markov state modeling
polyQ: Polygluatmine
RNP: ribonucleoprotein
RRM: RNA recognition motif
TICA: time-lagged independent component analysis

## Supporting Information

**Table S1:** Stationary probabilities (*π*), free energies (*G*), and average protein heavy-atom root-mean-square deviations (RMSDs) of the metastable states, computed after alignment to the crystal structure.

| State | $\pi$ (%) | $G$ (kcal/mol) | RMSD (nm) |
| --- | --- | --- | --- |
| S1 | $2.38 \pm 0.78$ | $1.946 \pm 0.196$ | $0.429 \pm 0.018$ |
| S2 | $5.10 \pm 0.90$ | $1.492 \pm 0.105$ | $0.401 \pm 0.019$ |
| S3 | $2.02 \pm 2.14$ | $2.045 \pm 0.633$ | $0.430 \pm 0.018$ |
| S4 | $3.78 \pm 2.38$ | $1.670 \pm 0.375$ | $0.453 \pm 0.017$ |
| S5 | $24.4 \pm 2.965$ | $0.558 \pm 0.072$ | $0.394 \pm 0.019$ |
| S6 | $62.2 \pm 4.826$ | $0.000 \pm 0.046$ | $0.395 \pm 0.014$ |

**Figure S1:**
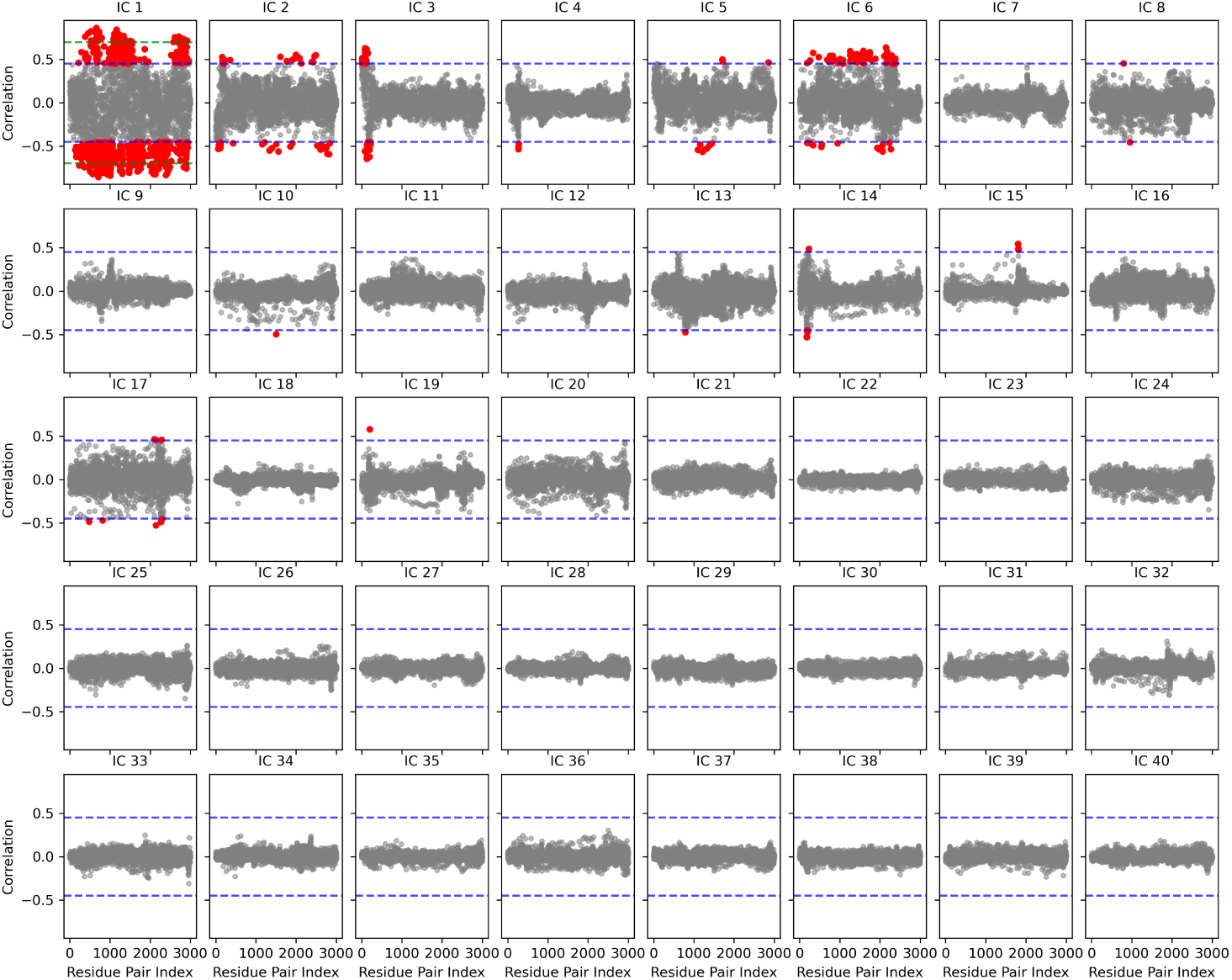
Feature–TICA correlations of residue pairs for the first 40 independent components. Correlation values above 0.45 are shown in red, and values below 0.45 are shown in gray. The horizontal blue line indicates the 0.45 threshold, and the green line indicates 0.70.

**Figure S2:**
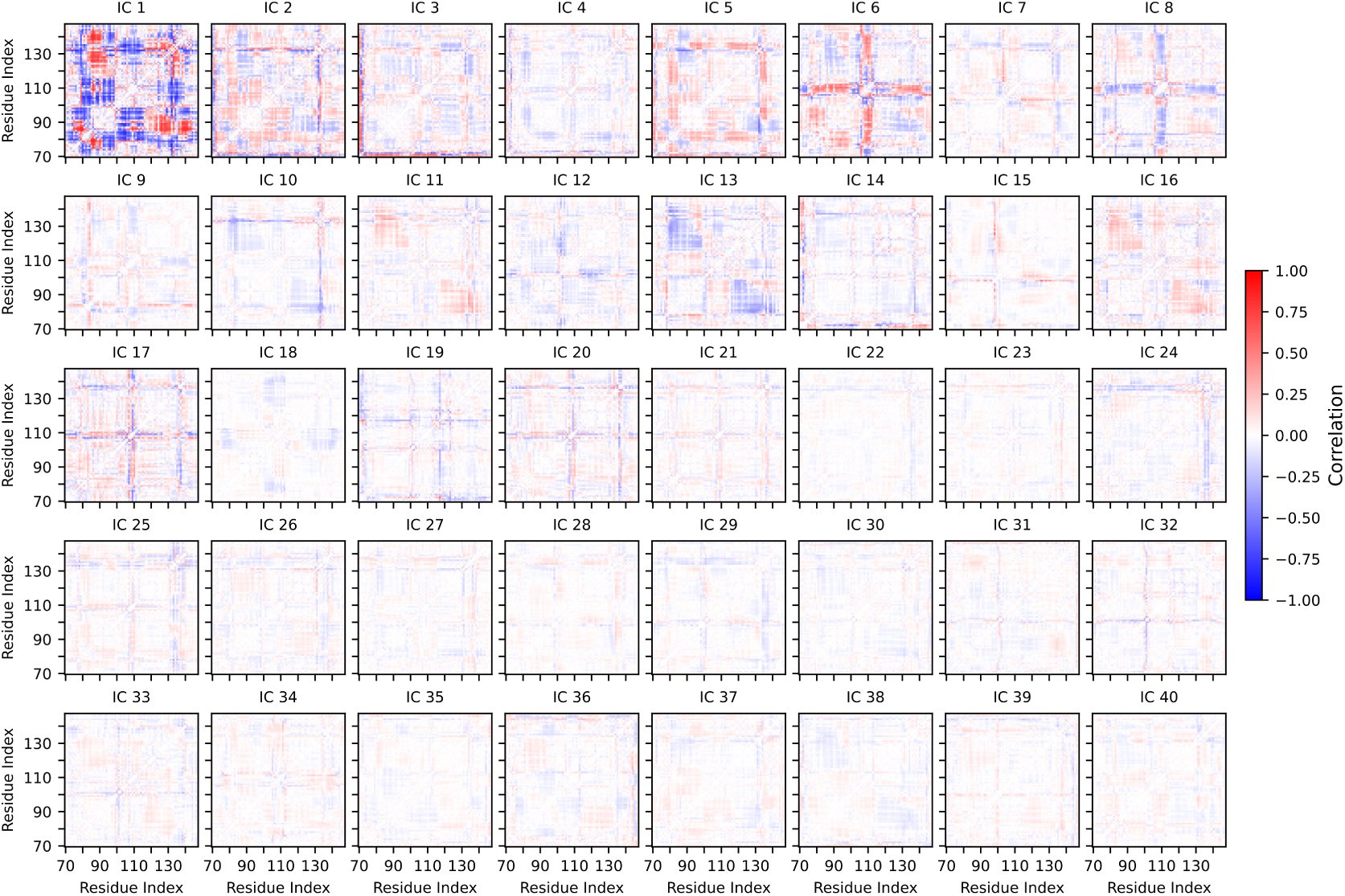
Feature-TICA correlations of residue pairs for the first 40 independent components, shown as a matrix. The color bar indicates the correlation values.

**Figure S3:**
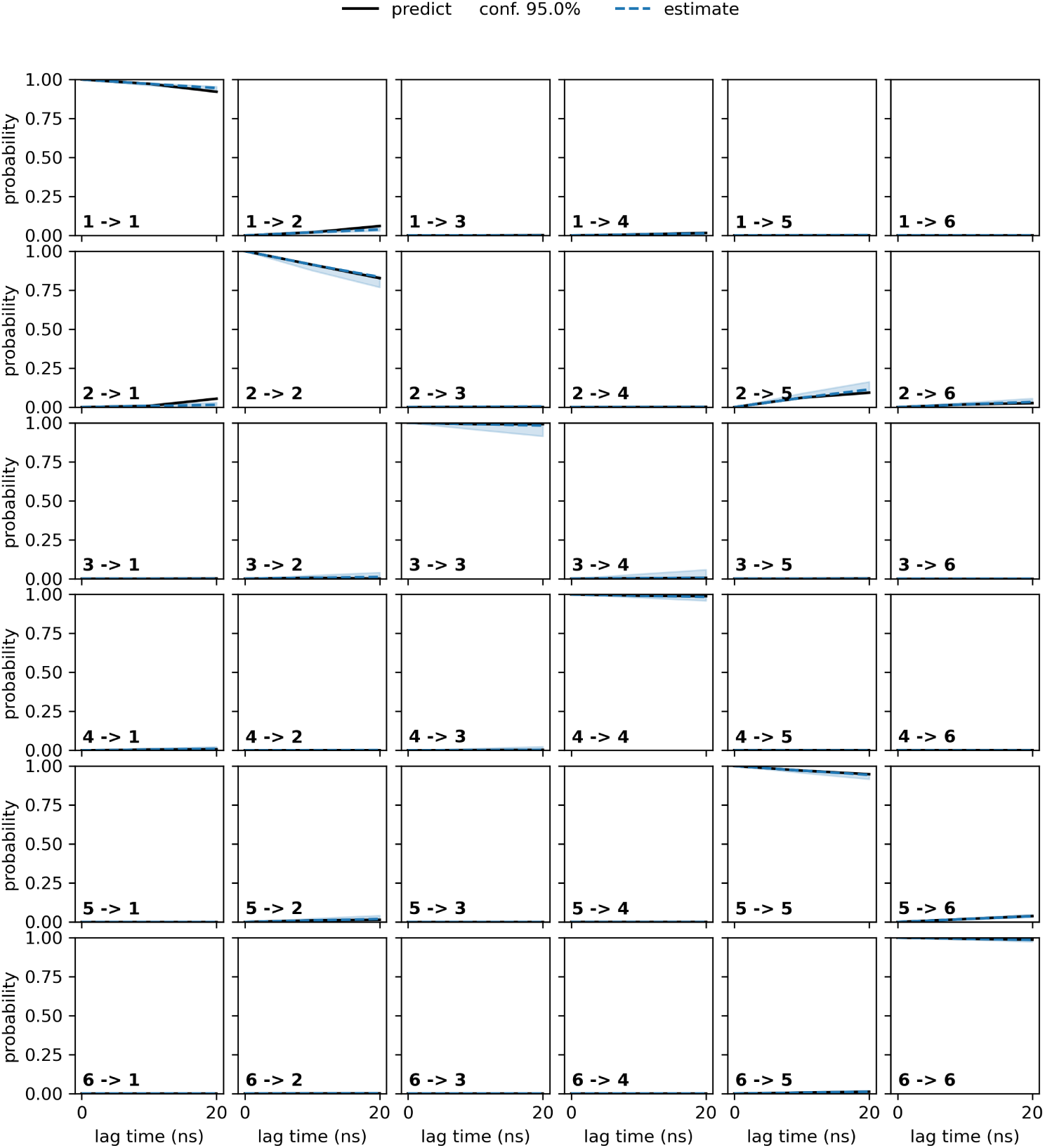
Chapman-Kolmogorov test for BHMSM validation at a lag time of 10 ns with six states. The estimated models are represented by blue solid lines, while the predictions are shown as black dashed lines. The shaded regions represent the 95% confidence intervals of the estimated models.

**Figure S4:**
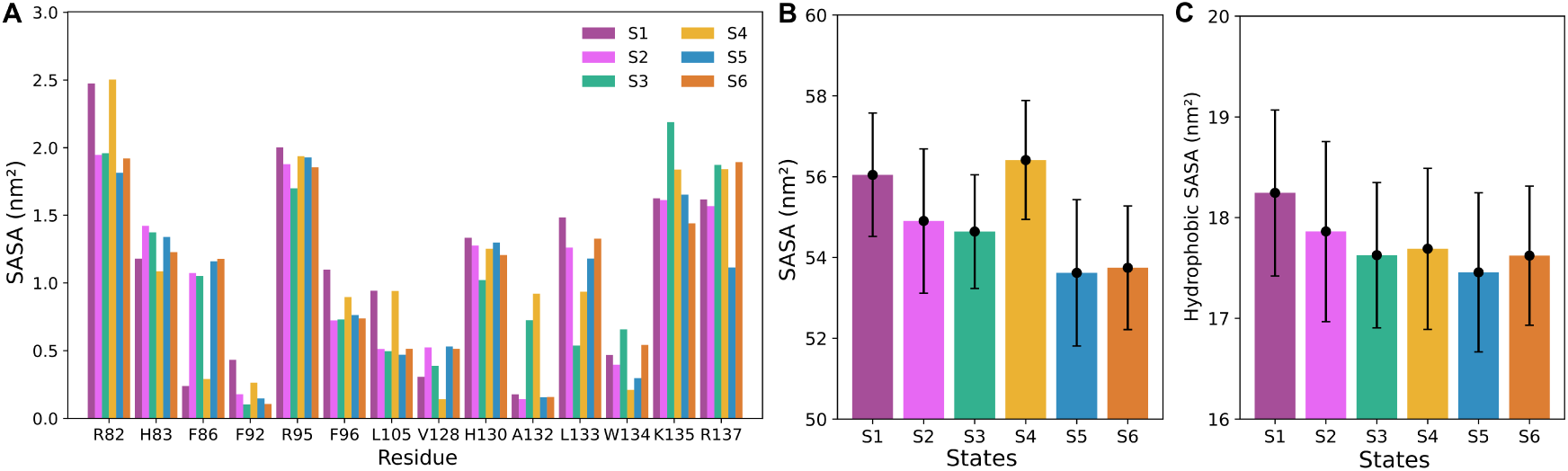
(A) Average solvent-accessible surface area (SASA) of residues showing significant differences (> 0.3 nm^2^) across the metastable states. (B) Average total SASA of the protein in each metastable state. (C) Average SASA of the protein hydrophobic residues in each metastable state. Error bars represent standard deviations.

**Figure S5:**
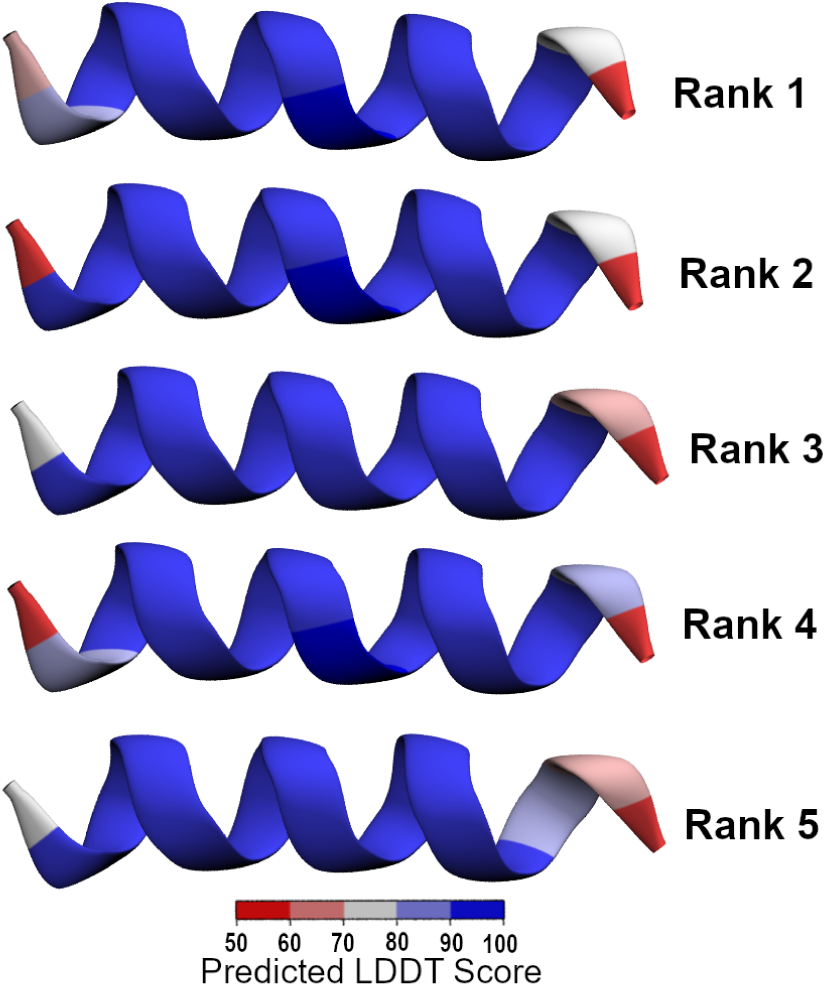
Predicted AlphaFold2 structures for residues 119-134, encompassing the *α*2-helix and loop5. Structures are colored according to their predicted LDDT (local distance difference test) scores (red to blue), where scores > 90 indicate high confidence, and scores between 70-90 indicate regions expected to be modeled accurately.

